# Human frontal eye field and eyelid motor area revisited with electrical cortical stimulation and electrode co-registration

**DOI:** 10.64898/2026.03.12.711460

**Authors:** Tomoyuki Fumuro, Juan C Bulacio, William E Bingaman, Akio Ikeda, Hiroshi Shibasaki, Hans O Lüders, Dileep R Nair, Riki Matsumoto

**Author notes:** **Corresponding author:** Riki Matsumoto, M.D., Ph.D., Department of Neurology, Graduate School of Medicine, Kyoto University, 54 Kawara-cho, Shogoin, Sakyo-ku, Kyoto City, Kyoto Prefecture, 606-8507, JAPAN. Deceased author: Died June 9, 2022 (Hiroshi Shibasaki).

## Abstract

We investigated the anatomical localization of the frontal eye field (FEF) and its relationship to the eyelid motor area (EMA) and precentral motor cortex. We performed functional mapping using electrical cortical stimulation (ECS) and correlated electrode position by non-linear co-registration techniques using postoperative MRI. We studied 22 patients who underwent chronic implantation of subdural electrodes for epilepsy surgery. Eye movements were elicited at 52 electrodes overall. The majority of the movements were conjugated, saccadic eye deviation contralateral to the side of ECS. Head turning and non-saccadic eye deviation more frequently occurred in the vicinity of the precentral sulcus. Anatomically, FEF was located at Brodmann’s area 6 in the most-caudal region of the middle frontal gyrus and in the adjacent part of the superior frontal sulcus and precentral sulcus. Functionally, FEF was situated at the level of the hand motor area, more dorsal than was described in Penfield’s motor homunculus. The FEF is situated anteriorly from the precentral motor cortex. The EMA was situated within the precentral motor cortex, partially overlapping with but distinctly ventral and caudal to FEF, and dorsal to the lower face motor area. A standardized map of the FEF and precentral motor homunculus is provided as a reference for human system neuroscience research.

## 1. Introduction

In the field of clinical functional mapping for patients who undergo tumor or epilepsy surgery, direct electrical cortical stimulation (ECS) has been a gold standard measure for functional cortical mapping since the early 20^th^ century (Penfield & Jasper, 1954). Transient functional alternation induced by this method can provide patient-specific identification of the motor homunculus or the precentral motor cortex. Accurate identification of oculomotor-related regions, including the frontal eye field (FEF) and the eyelid motor area (EMA), is clinically important for preserving motor function during resective surgery involving the perirolandic region.

The FEF, which is located at the rostral bank of the arcuate sulcus in Brodmann’s area (BA) 8 in the monkey (Bruce & Goldberg, 1985; Bruce, Goldberg, Bushnell, & Stanton, 1985), has been studied in this context in humans. In Penfield’s era (1954), by means of intraoperative ECS during awake craniotomy, FEF sites were considered mainly anterior to the face motor area in BA 8. In the late 1980s, Lüders et al. (1987) established the functional mapping with ECS using chronically implanted subdural electrodes for presurgical evaluations, which has enabled the detailed investigation of normal cortical functions by carefully monitoring the absence of afterdischarges on the electrocorticogram (ECoG). Recent ECS studies in the chronic setting have suggested that FEF sites are located anterior to the hand motor area and posterior to the silent area (Godoy, Lüders, Dinner, Morris, & Wyllie, 1990; Blanke et al., 2000). Dorsoventrally, FEF sites mainly lie ventral to the upper extremity and dorsal to the face motor area (Godoy et al., 1990).

Advancements in functional neuroimaging techniques such as PET activation study using a H_2_^15^O tracer or functional MRI have enabled scientists to study the location of FEF in healthy subjects. Human FEF was located at the caudal middle frontal gyrus (MFG), specifically at the junction of the precentral sulcus (PrCS) and the superior frontal sulcus (SFS) in BA 6 (Petit, Clark, Ingeholm, & Haxby, 1997; Luna et al., 1998; Berman et al., 1999; Rosano et al., 2002). The localization of the FEF was more dorsal than that reported by Penfield et al. (1954). Only a few ECS studies have combined anatomical neuroimaging for precise localization of the FEF. To date, such attempts have only been performed in small patient populations with subdural electrodes (Blanke et al., 2000) or using depth electrodes (Lobel et al., 2001). Over the past decade, depth electrode-based ECS studies have emerged as the standard approach for functional mapping, driven by the widespread adoption of stereo-EEG (SEEG) procedures as the primary method for invasive presurgical evaluation of intractable focal epilepsy in most epilepsy centers. Though depth electrodes are suitable for investigating functions in the cerebral sulcus, they cannot detect the spatial distribution of a particular function on the brain’s surface. On the other hand, grid or strip subdural electrodes have an advantage in investigating cortical functions surrounding the seizure-onset zone in a two-dimensional way over the surface of the brain, albeit the investigation area is limited to the crown part of the cerebral gyri. Given the widespread adoption of SEEG for presurgical evaluations of intractable focal epilepsy, a systemic prospective ECS mapping study using subdural electrodes is not anticipated globally. Therefore, we opted to retrospectively include patients who underwent presurgical evaluations with chronically implanted subdural electrodes dating back to approximately the year 2000, aiming to better characterize the anatomical and functional properties of the FEF. These patients underwent ECS in the epilepsy monitoring unit under awake conditions with detailed video analysis for eye and other body movements. These patients also underwent MRI scans after the implantation of subdural grids, which enabled researchers to define anatomical locations of electrodes in relation to cerebral sulci and perform co-registration into the Montreal Neurological Institute (MNI) standard space. We feel that this methodology offers a higher level of precision compared to using CT scans performed after implantation (Matsumoto et al., 2004). This study aimed to delineate the anatomo-functional characteristics of the FEF; i.e., the relationship with cerebral sulci and other portions of the motor homunculus, and carry out detailed analyses of the oculomotor response and accompanying head turning.

The face portion of the motor homunculus has been extensively studied with ECS for the lower face, such as the mouth and tongue; however, there is a paucity of findings for the upper face. Penfield et al. reported the variable location of the EMA along the precentral gyrus and adjacent frontal convolution, partly because of the location of electrodes were identified visually (Penfield & Boldrey, 1937; Rasmussen & Penfield, 1948). Moreover, only a few fMRI studies are available for voluntarily blink movements in healthy subjects (Kato & Miyauchi, 2003a; Bristow, Haynes, Sylvester, Frith, & Rees, 2005; Hanakawa, Dimyan, & Hallett, 2008; van Koningsbruggen, Peelen, Davies, & Rafal, 2012). Our second objective is to characterize the anatomical and functional properties of the EMA. To our knowledge, no studies have investigated EMA characteristics by combining ECS and precise anatomical imaging. In this study, we identified the location of the EMA and found evidence that different regions are responsible for eyelid motor and oculomotor response.

In addition, we also attempted to generate a standardized map of the FEF, EMA, and other motor areas in the precentral motor cortex by co-registering the data obtained in individual brains into the MNI standard space as a reference for human system neuroscience.

## 2. Materials and Methods

### 2.1 Subjects

Between January 1997 and March 2002, 101 medically intractable patients were evaluated for surgery with chronic implantation of subdural electrodes. The surgical technique and details of the subdural electrode arrays are discussed elsewhere (Godoy et al., 1990). The implanted electrodes, which measured 3.97 mm in diameter with a center-to-center inter-electrode distance of 1 cm, were made of platinum (custom-made in the Cleveland Clinic, OH, U.S.A.).

Sixty-eight patients had subdural electrode coverage over the lateral frontoparietal area, including the premotor or precentral area. An 8 × 8, 4 × 11, or 8 × 5 electrode grid or grids with combinations were implanted in the perirolandic region for functional mapping. In 58 patients, ECS was performed for the localization of eloquent cortices in and around the perirolandic region. In 35 (59%) patients, eye movements were elicited by ECS. Three patients had a lesion or epileptic focus in the premotor or precentral area and were excluded from the analysis. Twenty-two patients were finally recruited into the present study, with video recordings of cortical stimulation available for review. The present study was approved by the Institutional Review Board Committee of the Cleveland Clinic Foundation (IRB #4513), and all data were obtained at the Cleveland Clinic as part of the presurgical evaluation for epilepsy surgery.

### 2.2 Functional cortical mapping via ECS

ECS was performed for functional mapping in all subjects as a part of the presurgical evaluation. The method has been reported in detail elsewhere (Lüders et al., 1987; Matsumoto et al., 2007). The procedure was performed outside the operating room with the patients fully awake, sitting comfortably in bed, with their heads unstrained. Repetitive square wave electric currents of alternating polarity with a pulse width of 0.3 ms and a frequency of 50 Hz were delivered to each subdural electrode (Grass S-88 and SUI-7, Asro-Med Inc., R.I., U.S.A.). Train duration was set to 5 s. In each patient, the subdural reference electrode for the pseudomonopolar ECS stimulation was selected based on absence of interictal or ictal epileptiform discharges and absence of positive response. This electrode was used as the reference for stimulating all the other electrodes. Throughout stimulation, ECoG were monitored continuously to observe any induced afterdischarges or EEG seizure patterns. Stimulus current was started at 1 mA and increased gradually (in 0.5 mA to 1 mA steps) until (1) a positive response was observed, (2) the maximum intensity of 15 mA was reached, or (3) afterdischarges were elicited with an intensity of <15 mA. An FEF electrode was defined as the electrode at which eye movements were observed without afterdischarges being induced by stimulation. Once an oculomotor response was obtained, subsequent stimulation was applied while the patient fixated on an object in the central position with his/her eyes and head aligned. Only positive/negative responses reproducible more than twice were used for eye and other movement analyses.

### 2.3 Eye movement analysis

All videotapes were reviewed independently by two authors (RM, DN). We determined the duration, type, and range of movements of the elicited eye deviation, as well as the presence of the associated head version and face motor responses.

Eye deviation was considered as full-range deviation when the sclerocorneal border reached the outer canthus of the eye ipsilateral to the deviation. When a clear-cut jerky eye movement was identified, it was labeled “saccadic.” If the deviation appeared to be continuous, without saccades, it was classified as “non-saccadic.” When the observers did not agree on the type of movement observed, it was labeled “inconclusive.”

The presence of vertical components of eye movements was also analyzed. Regarding the direction of the eye movement, we defined it as “horizontal” when the eye moved on a straight line between the inner and outer canthus and defined everything else as “oblique.”

Associated head version was considered to be present when a clearly defined involuntary head turning (HT) occurred either during or immediately after the eye movements. HT was subdivided into two categories: HT that started before the full-range eye deviation (“HT before-end”) and HT that followed the full eye deviation; i.e., HT starting after full-range eye deviation was achieved (“HT after-end”).

### 2.4 Anatomical-functional localization mapping of the FEF

The anatomical location of each subdural electrode was identified with three-dimensional (3D) MRI images taken after the implantation of subdural grids. Details of this methodology have been described elsewhere (Matsumoto et al., 2004). T1-weighted MRI images were acquired and combined to form a single 3D volume (1.5 T, MPRAGE sequence). The location of each electrode on a subdural grid was identified on the two-dimensional (2D) MR images (Fig. 1A) using its void signal due to the property of the platinum alloy. The surface of the brain was reconstructed using an in-house computer program that interactively renders brain volume and surface locations of the subdural electrodes, using a Silicon Graphics computer (Mountain View, CA, USA). To represent each electrode in 2D position coordinates, a sphere was surface-rendered at the centroid of each identified electrode and overlaid on the volume-rendered image (Fig. 1A). To compensate for artifacts arising from subdural grids on the surface of the brain, the volume-rendered MRI was chamfered by 4–5 mm from the cortical surface to locate the sulci beneath the grids (Fig. 1B). Sulcal patterns in the lateral frontal region were identified based on the atlas in a previous study (Ono, Kubik, & Abernathey, 1990) (Fig. 1C). Because the intersection of the SFS and the PrCS is a constant landmark of the adult human brain (Ono et al., 1990) and the location of the FEF has been strongly associated with these two sulci in previous imaging studies (Paus, 1996), the location of each FEF electrode was identified with reference to the SFS and PrCS (Fig. 1D). Straight lines were drawn as virtual sulci to best represent the portion of the SFS and PrCS close to their junction, and distances from the SFS and PrCS were measured by drawing a straight line to each virtual sulcus (Fig. 1E). If the PrCS was divided into the superior and inferior rami, the distance from the PrCS was measured from a virtual PrCS connecting two junctions between the superior ramus and the SFS and between the inferior ramus and the inferior frontal sulcus (IFS). When the ramus did not actually connect with the SFS or IFS due to its variation, the virtual junction was defined as the meeting point of each ramus and the extended lines of the SFS or IFS. Regarding the axes of coordinates, we defined the 2D dorso-ventral axis as corresponding to the PrCS, and the 2D rostro-caudal axis as corresponding to the SFS. The location of FEF electrodes was plotted in a coordinate system where the 2D dorso-ventral axis showed the distance of the SFS and 2D rostro-caudal axis from the PrCS with an origin representing the meeting point of the SFS and PrCS (Figs. 1E, F). Thus, the quadrant with 2D dorso-ventral axis > 0, 2D rostro-caudal axis > 0 corresponds to the MFG or potentially the inferior frontal gyrus (IFG), and that with a 2D dorso-ventral axis < 0, 2D rostro-caudal axis > 0 to the superior frontal gyrus (SFG). Similarly, the precentral gyrus (PrCG) is represented by the coordinate of the 2D rostro-caudal axis < 0 (Figs. 1E, F). For precise localization of FEF electrodes over these sulci, each electrode was analyzed in relation to the real sulci in individual chamfered 3D MRI.

**Figure 1.**
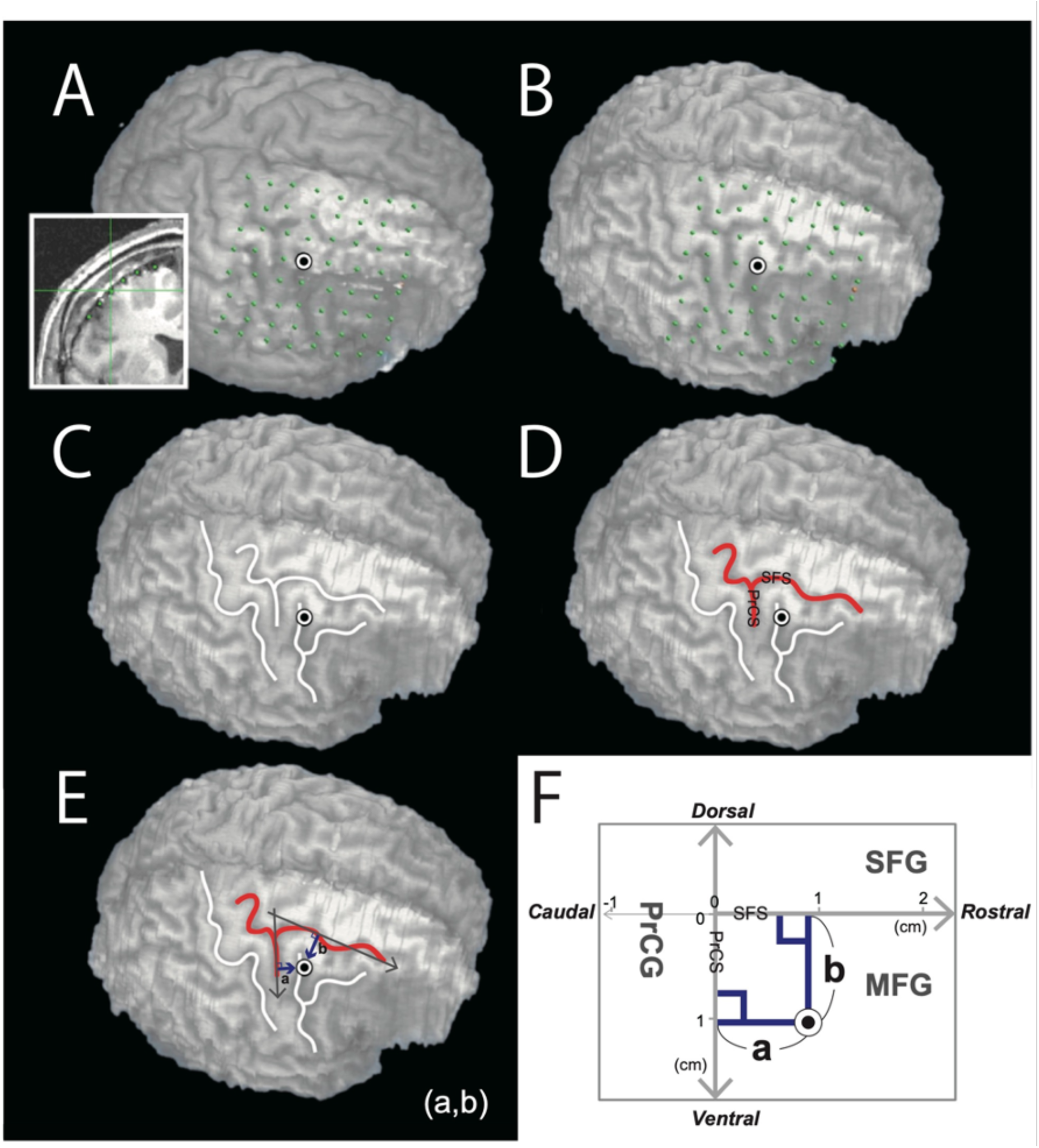
Co-registration of subdural electrodes to T1 volumetry taken after electrode implantation. (A) The location of each electrode on a subdural grid was identified in 2D MR images using its void signal due to the property of platinum alloy. A sphere was surface-rendered at the centroid of each identified electrode and overlaid on the volume-rendered image. (B) The volume-rendered MRI was chamfered by 4–5 mm from the cortical surface to locate the sulci beneath the grids. (C) The sulcal patterns in the lateral frontal region were identified. (D) The location of each FEF electrode was identified with reference to the SFS and PrCS. (E, F) The location of FEF electrodes was plotted in a coordinate system where the 2D dorso-ventral axis shows the distance from the SFS and the 2D rostro-caudal axis from the PrCS with an origin representing the meeting point of the SFS and PrCS.

We divided the electrodes into three groups based on their electrical current threshold (low: ∼5 mA, middle: 5.5–10 mA, and high: 10.5–15 mA) for eliciting eye movements. We performed the one-way analysis of variance (ANOVA) to determine whether the distances from the intersection of the SFS and PrCS differed significantly or not between the three threshold groups.

The distribution of the FEF in each subject was also investigated. For subjects with more than two FEF electrodes, the center of the FEF was calculated individually by measuring the mean coordinates in 2D dorso-ventral and 2D rostro-caudal axes in each subject.

The variance of the distribution of the FEF was explored along the 2D dorso-ventral and 2D rostro-caudal axes. To determine whether the degree of variance differed between the two axes, the coordinates of the FEF were compared. Levene’s test was used for inter-subject analysis. For subjects with more than two FEF electrodes, the maximum distance between electrodes was measured in each subject in the 2D dorso-ventral and 2D rostro-caudal axes, respectively. To compare their maximum distance between the two axes, the Wilcoxon signed-rank test was performed for within-subject analysis.

To evaluate the relationship between the coordinates of the two axes of FEF electrodes, i.e., to study whether the FEF was significantly shifted caudally as it was located more ventrally, Spearman’s rank correlation test was performed using the aforementioned coordinates for all FEF electrodes. To evaluate this shift on an individual basis, for subjects with more than two FEF electrodes, a linear regression line was drawn for these electrodes in each subject based on a least square method, to explore the coordinates in the 2D rostro-caudal axis on the 2D dorso-ventral axis. The sign test was used to identify whether the FEF was significantly shifted caudally as it was located more ventrally in each subject.

To evaluate the distributional difference between saccadic and non-saccadic eye movements, i.e. whether the non-saccadic electrodes were significantly proximal to the PrCS compared to saccadic electrodes, the unpaired *t*-test was performed using the distance of each electrode from the PrCS.

The direction of eye movement was similarly analyzed. The unpaired *t*-test was performed to compare their distance from each electrode to the PrCS between the oblique and the horizontal electrodes.

Regarding the HT, all FEF electrodes were classified into three groups: eye deviation without HT, HT before-end, and HT after-end. To determine whether there was a significant difference in the number of the electrodes gathered over PrCS between those 3 groups, Fisher’s exact test, residual analysis. In addition, the multivariate analysis of variance (MANOVA) was performed to examine whether there was a significant difference in electrode coordinates, i.e., 2D dorso-ventral and rostro-caudal axes, respectively, among these three groups.

### 2.5 Definition of the motor homunculus

Positive motor areas were defined based on the presence of muscle contraction with ECS occurring at the contralateral body with the expected somatotopy. Positive motor areas were divided into eight groups based on the segment involved: in the lower extremity (LE) (thigh, leg, or foot), arm, forearm, hand, FEF, EMA, lower face, or tongue. The methodology of eye movement analysis was described above. The eyelid motor responses were analyzed, if present alone or in combination with eye movements. Eyelid motor response was characterized using (1) side of contraction (bilaterally or unilaterally), (2) mode of contraction (tonic or clonic), and (3) presence of a concomitant positive motor response (the lower face motor response (i.e., the tongue or mouth) or oculomotor response). Only responses that the two authors (RM, DN) agreed on were accepted. Positive motor areas in the upper extremity were defined as follows: contraction of the intrinsic muscle was defined as the hand motor area, wrist movement was defined as the forearm motor area, and elbow/shoulder movement was defined as the arm motor area. The negative motor area (NMA) was defined as the cortical region, the stimulation of which elicited the inability to conduct voluntary movement in the absence of the positive motor phenomenon (Matsumoto et al., 2007). The language area was defined as the cortical region, which when stimulated showed impairments in reading aloud in the absence of a negative motor response.

### 2.6 Comprehensive anatomical-functional mapping in the MNI standard space

Individual electrode coordinates were transferred to the MNI standard space (ICBM-152) to evaluate the distribution of all ECS motor responses in the precentral motor cortex for all patients. Electrodes identified in the T1 volume acquisition (1.5 T, MPRAGE) obtained after grid implantation were non-linearly co-registered to the T1 volume acquisition before implantation (1.5T, MPRAGE). These locations were then converted to the MNI standard space using FNIRT in the software FSL (www.fmrib.ox.ac.uk/fsl/fnirt/). This method has been reported elsewhere for the standardization of electrode locations (Matsumoto et al., 2011). The electrode MNI coordinates were transferred into Talairach coordinates (Talairach, Tournoux, & Rayport, 1988) using BrainMap GingerALE (http://www.brainmap.org). BA of each coordinate was identified using Talairach Client software (http://www.talairach.org/client.html). To analyze relative positions among electrode coordinates of the positive/NMAs, coordinates in the left hemisphere were all flipped into the right hemisphere in further analysis. All positive/negative motor electrodes revealed by cortical stimulation were plotted together in the MNI standard space to investigate the distribution with respect to each function. To represent each electrode, a sphere with a radius of 2 mm was surface-rendered at each electrode coordinate. The mean coordinate of each positive/negative motor area was also represented as a sphere with a radius of 5 mm. For comparison, coordinates of FEF, EMA, blink, and supplementary eye field from previous neuroimaging and/or cortical stimulation studies are also shown in the same figure.

### 2.7 Anatomical localization of the EMA reference to FEF in 2D and 3D analyses

Two sets of analyses were performed to delineate the location of the EMA with reference to the FEF. First, the positional relationship between each eyelid motor electrode and the PrCS was clarified. Using the Talairach Client software program, the numbers of eyelid motor electrodes that were located caudal or rostral to the PrCS were counted. The analysis was repeated for the FEF electrode. Fisher’s exact test was performed to determine whether the positional relationship between the EMA and PrCS was distinct from that between the FEF and PrCS.

Second, for subjects who showed separate eyelid motor and oculomotor responses, these locations were plotted in the same 2D coordinate to compare their anatomical location. For subjects with more than two FEF (or EMA) electrodes, the center of the FEF (or EMA) was calculated and plotted. The center coordinate was calculated individually by measuring the mean coordinates in the 2D dorso-ventral and 2D rostro-caudal axes in each subject. The MANOVA test was performed to determine whether the EMA is located distinctly from the FEF. In addition to the 2D analysis, The MANOVA test with the MNI standard space was also performed in three axes (X-axis [medial-lateral], Y-axis [rostro-caudal], and Z-axis [dorsal-ventral]) for the same purpose.

### 2.8 Surrounding functions of FEF and EMA in 2D analysis

To delineate the location of the FEF with reference to the precentral motor cortex, functions surrounding the FEF electrodes were investigated in the following axes: the rostro-caudal axis (parallel to the AC-PC line) and the dorso-ventral axis (parallel to the PrCS). Because the configuration of the grid was not usually aligned to the AC-PC line, the function of the electrode that predominates in the four directions along two axes (rostral, caudal, dorsal, ventral) was considered as 100% for this purpose. If a single electrode did not predominate in one direction, that is, if two electrodes almost equally (e.g., 40% to 60%) occupied the adjacent region in a particular direction, the functions of two electrodes were taken separately as 50% of the function in this particular direction. Some FEF electrodes were located at the edge of the grid or formed clusters with other FEF electrodes. When an FEF electrode was located at the grid edge, surrounding functions in that direction was considered indeterminate. When FEF electrodes formed a cluster, an electrode adjacent to an existing FEF electrode was also classified as an FEF electrode. Only the presence of motor function or silent areas were analyzed in the electrodes surrounding the FEF. The existence ratio of the surrounding functions was analyzed in each direction. Functions surrounding the eyelid motor electrodes were also investigated in the same way.

### 2.9 3D analysis of anatomical localization of the EMA relative to other positive motor areas

ANOVA and Dunnett’s *post-hoc* tests were performed to delineate the location of the EMA relative to the other positive motor areas. We examined whether the EMA was located in a distinct distribution along the Z-axis from the other positive motor areas.

### 2.10 Statistics analyses

All statistical analyses in the present study were performed using a commercially available statistical software program (SPSS 25.0 J for Windows; SPSS, Chicago, IL, USA). In both MANOVA and ANOVA, Tukey’s HSD tests were used for *post-hoc* analyses of significant effects.

## 3. Results

### 3.1 Patient demographics

The patient group consisted of 10 men (45%) and 12 women (55%). Their age ranged from 11 years to 47 years (mean: 25.3). Nineteen patients were right-handed (86%) and three were left-handed (14%). In each patient, 40–104 subdural electrodes (mean: 74.2) covered the lateral frontoparietal area, including the premotor and/or precentral area. Fifteen patients (68%) had subdural grids implanted in the left hemisphere and seven (32%) had the grids implanted in the right hemisphere. After presurgical evaluation with subdural electrodes, patients were diagnosed to have frontal lobe epilepsy (14 patients), temporal lobe epilepsy (four patients), or parietal lobe epilepsy (four patients), and a majority of the patients (21 patients) underwent surgical resection.

Table 1 shows brain functions and number of electrodes, patients, BA, and MNI coordinates. Every averaged coordinate, including LE, arm, forearm, hand, FEF, EMA, lower face, and tongue were located at BA 6. In addition to the 52 FEF electrodes, head turning was found at another electrode without oculomotor response. All 52 FEF and 14 EMA coordinates in the MNI standard space are shown in Supplements 1 and 2, respectively.

**Table 1.**
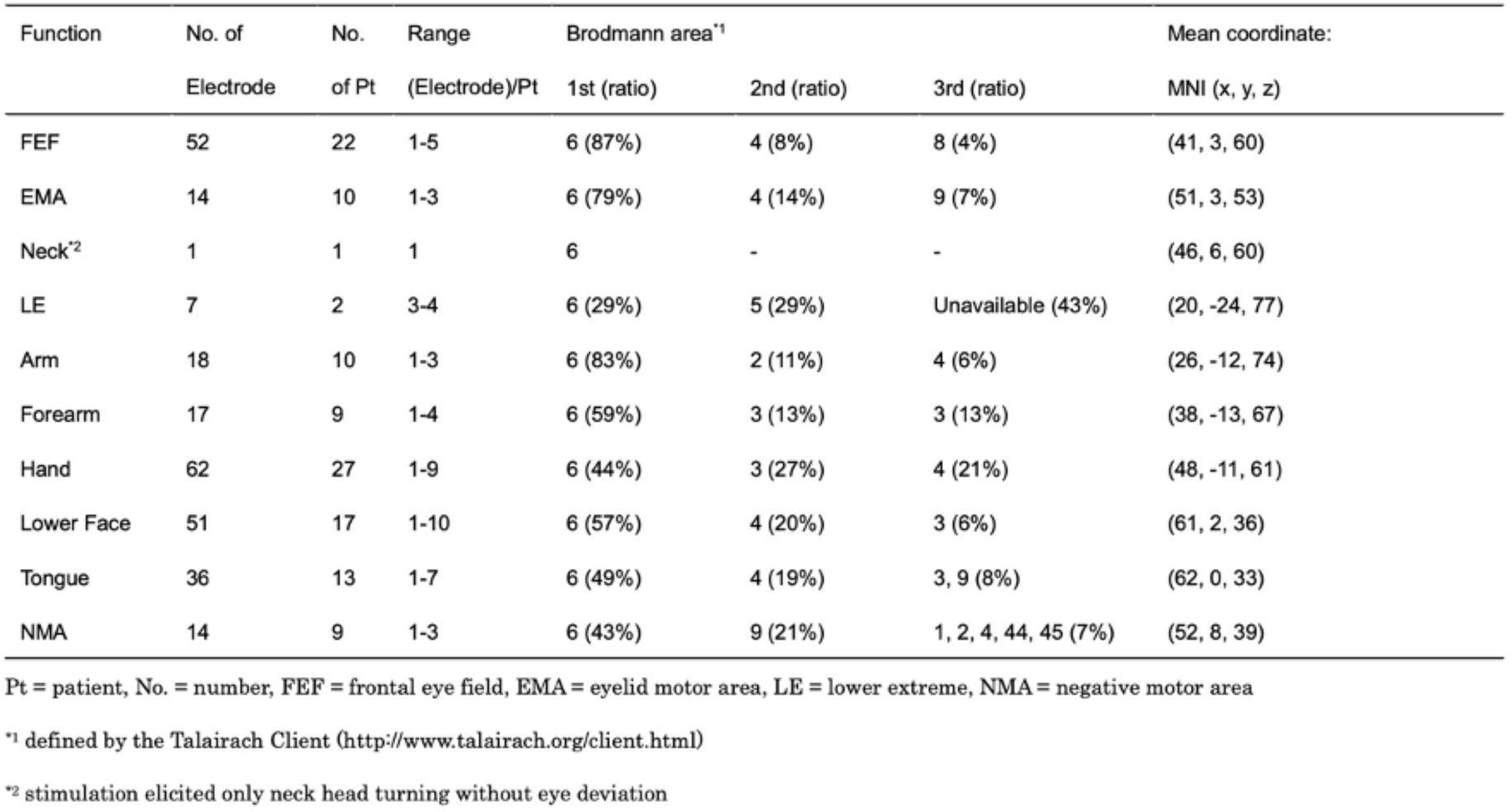
Functions elicited by cortical stimulation.

| Function | No. of<br>Electrode | No.<br>of Pt | Range<br>(Electrode)/Pt | Brodmann area<sup>1</sup> |  |  | Mean coordinate:<br>MNI (x, y, z) |
| --- | --- | --- | --- | --- | --- | --- | --- |
|  |  |  |  | 1st (ratio) | 2nd (ratio) | 3rd (ratio) |  |
| FEF | 52 | 22 | 1-5 | 6 (87%) | 4 (8%) | 8 (4%) | (41, 3, 60) |
| EMA | 14 | 10 | 1-3 | 6 (79%) | 4 (14%) | 9 (7%) | (51, 3, 53) |
| Neck<sup>2</sup> | 1 | 1 | 1 | 6 | - | - | (46, 6, 60) |
| LE | 7 | 2 | 3-4 | 6 (29%) | 5 (29%) | Unavailable (43%) | (20, -24, 77) |
| Arm | 18 | 10 | 1-3 | 6 (83%) | 2 (11%) | 4 (6%) | (26, -12, 74) |
| Forearm | 17 | 9 | 1-4 | 6 (59%) | 3 (13%) | 3 (13%) | (38, -13, 67) |
| Hand | 62 | 27 | 1-9 | 6 (44%) | 3 (27%) | 4 (21%) | (48, -11, 61) |
| Lower Face | 51 | 17 | 1-10 | 6 (57%) | 4 (20%) | 3 (6%) | (61, 2, 36) |
| Tongue | 36 | 13 | 1-7 | 6 (49%) | 4 (19%) | 3, 9 (8%) | (62, 0, 33) |
| NMA | 14 | 9 | 1-3 | 6 (43%) | 9 (21%) | 1, 2, 4, 44, 45 (7%) | (52, 8, 39) |

Pt = patient, No. = number, FEF = frontal eye field, EMA = eyelid motor area, LE = lower extreme, NMA = negative motor area
<sup>1</sup> defined by the Talairach Client (http://www.talairach.org/client.html)
<sup>2</sup> stimulation elicited only neck head turning without eye deviation

**Supplementary 1.** Coordinates of 52 FEF electrodes in the MNI standard space.

| x | y | z | x | y | z |
| --- | --- | --- | --- | --- | --- |
| 22 | -24 | 78 | 38 | -8 | 68 |
| 22 | -8 | 76 | 38 | -2 | 66 |
| 24 | 6 | 70 | 38 | 0 | 64 |
| 24 | 10 | 70 | 40 | -10 | 68 |
| 26 | -2 | 72 | 40 | 2 | 62 |
| 26 | 6 | 70 | 42 | 0 | 62 |
| 28 | -4 | 70 | 42 | 8 | 60 |
| 30 | -8 | 70 | 42 | 10 | 60 |
| 30 | 6 | 68 | 42 | 10 | 62 |
| 32 | 0 | 68 | 44 | 6 | 60 |
| 34 | 12 | 66 | 44 | 14 | 56 |
| 34 | 20 | 62 | 46 | -4 | 60 |
| 36 | -4 | 66 | 46 | -2 | 62 |
| 36 | 4 | 64 | 46 | 0 | 58 |
| 36 | 4 | 64 | 46 | 0 | 58 |
| 36 | 8 | 64 | 46 | 0 | 58 |
| 36 | 8 | 66 | 46 | 0 | 58 |
| 36 | 10 | 64 | 46 | 2 | 56 |

| x | y | z |
| --- | --- | --- |
| 46 | 2 | 58 |
| 46 | 4 | 60 |
| 46 | 8 | 58 |
| 48 | -2 | 56 |
| 48 | 8 | 52 |
| 50 | 8 | 54 |
| 52 | 10 | 46 |
| 52 | 16 | 44 |
| 52 | 24 | 38 |
| 54 | -2 | 50 |
| 54 | 0 | 50 |
| 54 | 8 | 48 |
| 56 | -2 | 52 |
| 56 | 6 | 46 |
| 58 | 0 | 46 |
| 60 | -12 | 50 |

**Supplementary 2.** Coordinates of 14 eyelid motor area electrodes in the MNI standard space.

| x | y | z |
| --- | --- | --- |
| 34 | 12 | 66 |
| 46 | -4 | 62 |
| 46 | 0 | 58 |
| 46 | 0 | 60 |
| 46 | 4 | 60 |
| 52 | 14 | 48 |
| 54 | -2 | 50 |
| 54 | 0 | 50 |
| 54 | 2 | 52 |
| 54 | 2 | 54 |
| 54 | 4 | 50 |
| 54 | 16 | 34 |
| 58 | -4 | 50 |
| 62 | -8 | 42 |

### 3.2 Anatomical localization of the FEF in the individual space via ECS

A total of 52 FEF electrodes were identified by ECS in all the participants. The number of FEF electrodes per subject was 2.36 on average and ranged from one to four. All FEF electrodes were situated contiguously in all but two subjects. All FEF electrodes were located within 1.5 cm of the PrCS. FEF electrodes were located in the following regions in order of abundance: MFG (19 electrodes, 36.5%), PrCS (13.5, 26.0%), PrCG (9, 17.3%), SFG (6, 11.5%), and SFS (4.5, 8.7%). One electrode was located over the point of intersection of the SFS and PrCS. Regarding FEF electrodes, the majority were located in the dorso-caudal region of the MFG or the adjacent part of the PrCS (except for one electrode located over the PrCS dorsal to the SFS) and the SFS (36 electrodes, 69.2%). No FEF electrodes were located in the IFG.

The electric current required for eliciting eye movements was 9.4 mA on average, the range being 3–15 mA. Electrodes with a low-current-threshold group (∼5 mA) were clustered at the most dorso-caudal region of the MFG (gross map; Fig. 2a). The one-way ANOVA showed that the distance from the intersection of the SFS and PrCS differed significantly between the three groups (F (2, 49) = 3.840, p = 0.028). *Post-hoc* Tukey’s HSD test revealed a significantly shorter distance from the intersection to the low-current-threshold group (∼5 mA), than to both the middle- (p = 0.024) and high-current-threshold groups (p = 0.024).

**Figure 2.**
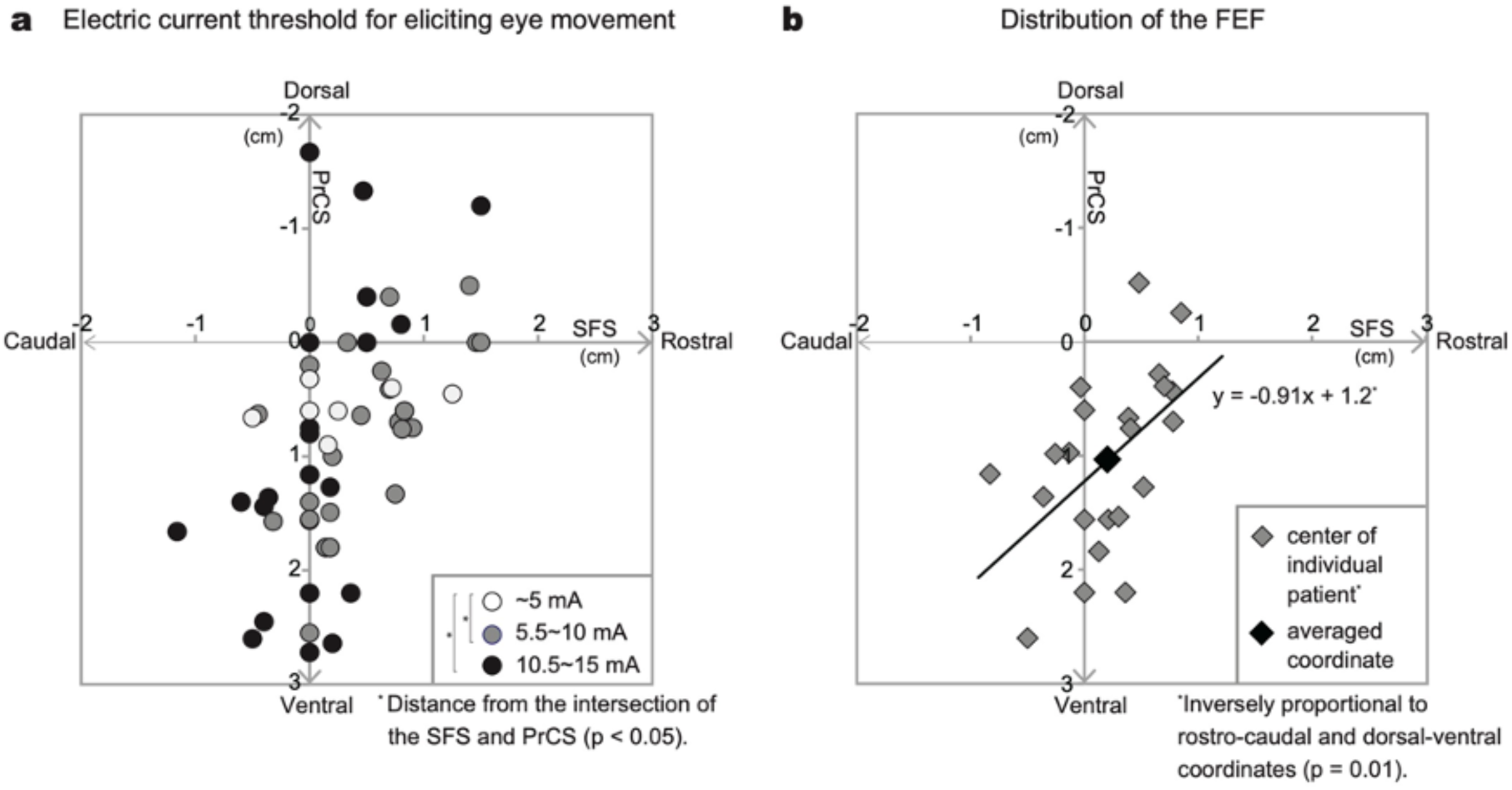
**(a)** FEF electrodes with corresponding intensity thresholds; low: 3–5 mA (white circle), middle: 5.5–10 mA (gray circle), high: 10.5–15 mA (black circle). **(b)** Center of FEF electrodes in individual patients (gray circle) and its averaged coordinate (black circle).

The center of the FEF in each subject is plotted in Fig. 2b, which shows the distribution of the FEF. The center of the FEF was located in the dorso-caudal region of the MFG or over the PrCS in the majority of the subjects (15 patients, 68.2%). The mean coordinate of the centers of FEF was located in the dorso-caudal region of the MFG ((2D dorso-ventral axis, 2D rostro-caudal axis) = (1.03, 0.20)) with a larger standard deviation along the 2D dorso-ventral axis (0.79) than along the 2D rostro-caudal axis (0.45). In the inter-subject analysis, the FEF had a significantly larger variance in the 2D dorso-ventral axis than in the 2D rostro-caudal axis (Levene’s test: p = 0.018). In the within-subject analysis of 15 subjects with more than two FEF electrodes, the maximal distance between FEF electrodes was also significantly longer in the 2D dorso-ventral axis than in the 2D rostro-caudal axis (mean ± standard deviation: 2D dorso-ventral axis 1.42 ± 0.45, 2D rostro-caudal axis 0.86 ± 0.49; Wilcoxon signed-rank test: p = 0.036). Thus, both inter- and within-subject analyses showed longer FEF distribution along the 2D dorso-ventral axis than the 2D rostro-caudal axis.

Spearman’s rank correlation analysis revealed that the coordinate in the 2D rostro-caudal axis decreased as that in the 2D dorso-ventral axis increased in both gross (52 electrodes (ρ = −0.54, p < 0.001) and center (22 patients with FEF electrodes: ρ = −0.54, p = 0.01) maps (Fig. 2b). In within-subject analysis of 15 subjects with more than two FEF electrodes, the mean gradient of a regression line was −1.45 with a range from −7.22 to 3.09. Thirteen subjects showed negative gradients among these subjects (sign test: p = 0.007), suggesting a caudal shift of the FEF electrodes in the ventral part of the FEF in each individual.

### 3.3 Eye movement characteristics of the FEF

As for the type of eye movements, saccadic movement was seen in 39 (73.6%) electrodes and non-saccadic movement in 15.1% (8 electrodes) (Fig. 3a). The type was inconclusive in four electrodes (7.5%). Full-range eye deviation was found in 37 (71.2%) electrodes. The averaged coordinates of non-saccadic electrodes were shifted caudally compared to those of saccadic electrodes (mean ± standard deviation of coordinates in the 2D rostro-caudal axis: non-saccadic 0.11 ± 0.68, saccadic 0.36 ± 0.59). In the 2D rostro-caudal axis, the saccadic electrodes varied between −1.16 and 1.5 from the PrCS. Non-saccadic electrodes were located within ± 0.5 cm from the PrCS. Upon comparing absolute values of the distance of each electrode from the PrCS between saccadic and non-saccadic electrodes, a significant difference in distribution was found between them (unpaired *t*-test: p = 0.019).

**Figure 3.**
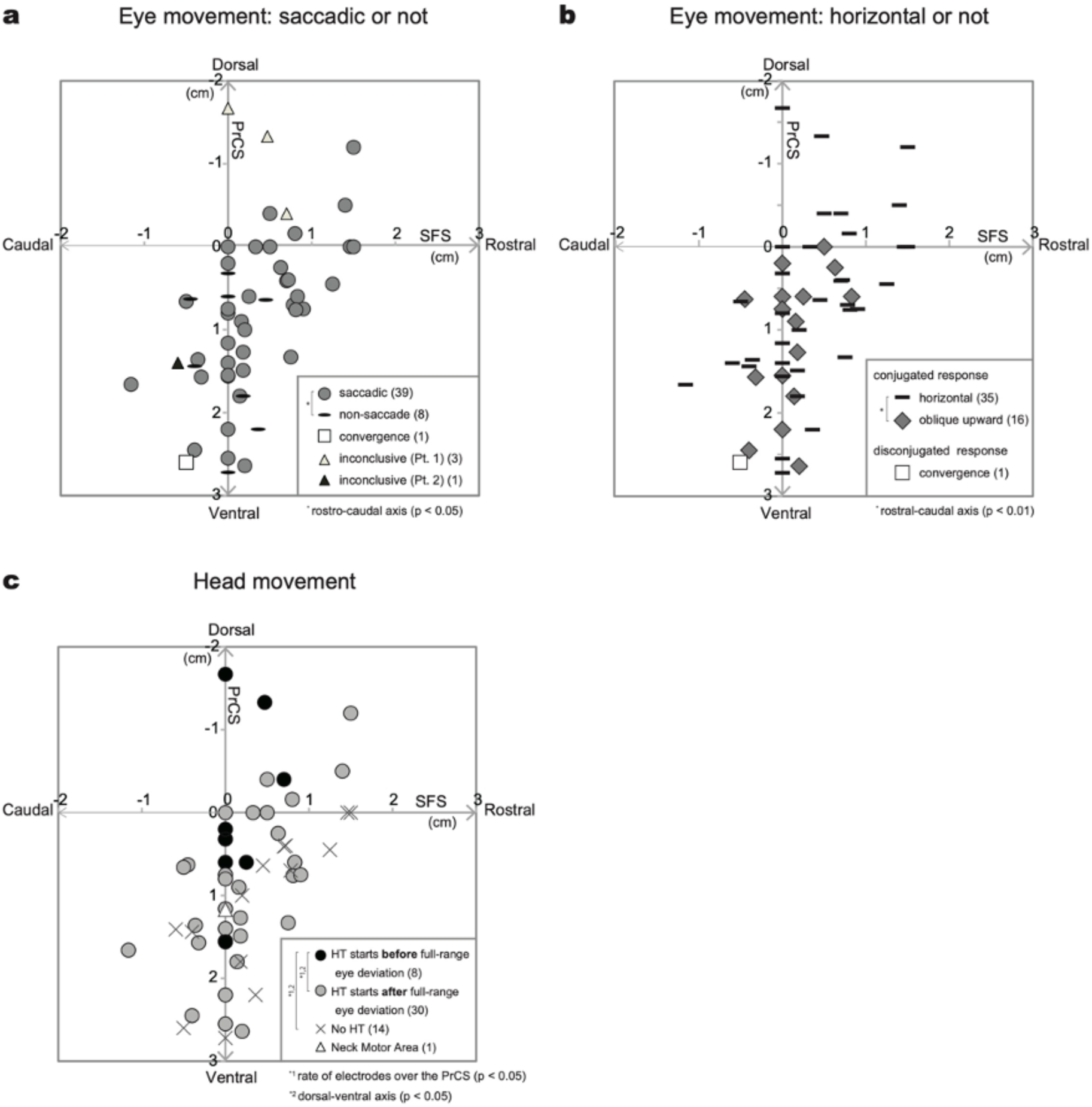
**(a)** The types of eye movements, saccadic and non-saccadic movements, are shown. Gray circle: saccadic movement, bar: non-saccadic movement, square: convergence movement. Inconclusive electrodes are shown in triangles. Triangles in white and black were derived from two patients. **(b)** The types of eye movement, conjugated and disconjugated, are shown. Bar: purely horizontal movement, gray argyle: oblique movement (up), square: convergence movement. **(c)** The characteristics of head turning are shown. The distribution of FEF electrodes with head version or head turning starting before full eye deviation (black circle) and after (gray circle). The crossing shows the electrodes without head deviation. The triangle shows head deviation alone without eye movement.

All the oculomotor responses observed were conjugated and contralateral to the side of cortical stimulation except for a disconjugated convergence response in one electrode (1.9%) in a single patient (Fig. 3b). Among the 51 FEF electrodes with conjugated responses directed contralaterally, 67.3% (35 electrodes) showed purely horizontal movement, whereas 30.8% (16 electrodes) had a vertical component that was always upward (oblique). The averaged coordinates of oblique electrodes were shifted caudally compared to those of horizontal electrodes (mean ± standard deviation of coordinate in 2D rostro-caudal axis: oblique 0.11 ± 0.35, horizontal 0.37 ± 0.63). In the 2D rostro-caudal axis, the horizontal electrodes varied between −1.16 and 1.5 from the PrCS. The oblique electrodes were located within −0.45 to 0.83 cm from the PrCS. Comparing the distance of each electrode from the PrCS between oblique and horizontal electrodes, there was a significant difference in distribution between them (unpaired *t*-test: p = 0.007).

HT was associated with conjugated eye deviation in 38 (73.1%) electrodes (Fig. 3c), and was always contralateral. Out of 38 FEF electrodes with HT, 30 electrodes were identified as HT after-end (i.e., HT starts after full-range eye deviation) while the remaining eight electrodes were HT before-end (i.e., HT starts before full-range eye deviation). HT never preceded the initiation of eye movements in these electrodes. In the 2D rostro-caudal axis, the rate of electrodes located over the PrCS (the coordinate of 2D rostro-caudal axis = 0) was 62.5% (5/8 electrodes) in HT before-end electrodes, 26.7% (8/30 electrodes) in HT after-end and 7% (1/14 electrodes) in eye deviation without HT. In terms of the number of electrodes over the PrCS, Fisher’s exact test showed a significant difference between the three groups (Fisher’s exact test: p = 0.020). The values of the expected number (EN) and adjusted standardized residuals (ASRs) indicated that the HT before-end electrodes were significantly aggregated into the PrCS (EN = 2.2, ASRs = 2.47, p = 0.014) compared to those of both HT after-end (EN = 8.1, ASRs = −0.04, p > 0.05) and eye deviation without HT (EN = 3.8, ASRs = −1.95, p > 0.05). Thus, the electrodes of eye deviation without HT rather tend to distribute outside the PrCS. The MANOVA was also performed on the electrode coordinates among these three groups, and a significant main effect was found (Wilk’s lambda = 0.714, F (4, 96) = 4.393, p = 0.003). Univariate analyses showed that the electrodes of HT before-end were located significantly dorsally compared with both HT after-end and eye deviation without HT only in the 2D dorso-ventral axis (mean ± standard deviation of coordinate in 2D dorso-ventral axis: HT before-end −0.02 ± 1.07, HT after-end 0.94 ± 0.94, eye deviation without HT 1.13 ± 0.92) (F (2, 49) = 4.017, p = 0.024. [*post-hoc* Tukey’s HSD] HT before-end–eye deviation without HT: p = 0.025. [*post-hoc* Tukey’s HSD] HT before-end–HT after-end: p = 0.039). There was no significant difference between them in the 2D rostro-caudal axis (p > 0.05).

As described above, in one electrode, a convergence response was observed (see a square electrode in Figs. 3a and 3b). This eye movement was the only oculomotor response in this patient. Neither HT nor a vertical eye movement was associated with the response. The convergence movement was smooth. The vergence angle was larger in the left eye, leading to a convergence response as if the patient had tried to focus on a near-space in the right hemifield, contralateral to the side of grid implantation. Tonic contractions were observed in both eyelids. The electrode was situated on the ventral margin of the superior ramus of the PrCS. The location of this electrode was identified in the following coordinate in the MNI standard space: (x, y, z) = (54, −2, 50).

In another electrode, cortical stimulation at an electrode situated caudally to the FEF electrode (see a triangular electrode in Fig. 3c) elicited HT alone. This subject was able to keep his eyes fixated on an object present in the center of the visual field during HT. Therefore, this area was thought to be the neck motor area. This neck motor area, after normalizing to the MNI space, corresponded to xyz coordinates of (46, 4, 60). In this subject, cortical stimulation to its rostral electrode elicited conjugate eye deviation with HT after-end. Other than HT, the only associated motor response was an eyelid motor response seen in four FEF electrodes across all subjects.

### 3.4 Characteristics of the EMA as observed via ECS

Eyelid motor area (EMA) was observed in 14 electrodes in 10 subjects. The number per subject ranged from one to three, the average being 1.4 electrodes. Electric current for eliciting eyelid motor response was 7.2 mA on average (range 4.5–15 mA). Eyelid closure was observed in eleven electrodes, while upper eyelid elevation was seen in three electrodes. Eyelid closure consisted of tonic (seven electrodes) or clonic (four electrodes) contraction and was seen bilaterally more on the side contralateral to cortical stimulation. Upper eyelid elevation consisted of unilateral tonic contractions and was seen contralateral to the side of stimulation. The lower face motor response (tongue or mouth) was associated with the eyelid response in five electrodes, and a simultaneous oculomotor response was seen in the other four electrodes.

### 3.5 Anatomical localization of the EMA relative to the FEF in 2D/3D analysis

Using the Talairach Client software, we defined the nearest gray matter from each eyelid motor electrode as being ten electrodes in the PrCG and four in the MFG. Similarly, the nearest gray matter from each FEF electrode was defined as being 26 electrodes in the MFG, 18 in the PrCG, 7 in the SFG, and 1 in the postcentral gyrus. Grouping locations of the electrodes into either caudal or rostral to the PrCS, the EMA had fewer electrodes that were rostral to the PrCS (4 of 14) than were located caudal to the PrCS (10 of 14). In contrast, the FEF had larger electrodes that were localized rather rostral to the PrCS (33 of 52) than caudal to the PrCS (19 of 52). Fisher’s exact test showed these differences between the EMA and the FEF to be statistically significant (p = 0.032).

The center of the EMA and FEF electrodes in each individual subject were plotted in the common coordinate. In total, there were six subjects who showed separate eyelid motor and oculomotor responses (Fig. 4). There was a significant difference in distribution between EMA and FEF (Wilk’s Lambda = 0.293, F (2, 9) = 10.88, p = 0.004) in both the 2D rostro-caudal axis (p = 0.001) and the 2D dorso-ventral axis (p = 0.006). The EMA was found to be more ventrally and caudally located than the FEF. In the MNI standard space as well, there was a significant difference in distribution between EMA and FEF (Wilk’s Lambda = 0.843, F (3, 62) = 3.863, p = 0.013) in X-axis (F (1, 64) = 11.77, p = 0.001) and Z-axis (F (1, 64) = 9.01, p = 0.004), but not in Y-axis (p > 0.05). This analysis confirmed that the EMA was identified to be located laterally and ventrally to the FEF.

**Figure 4.**
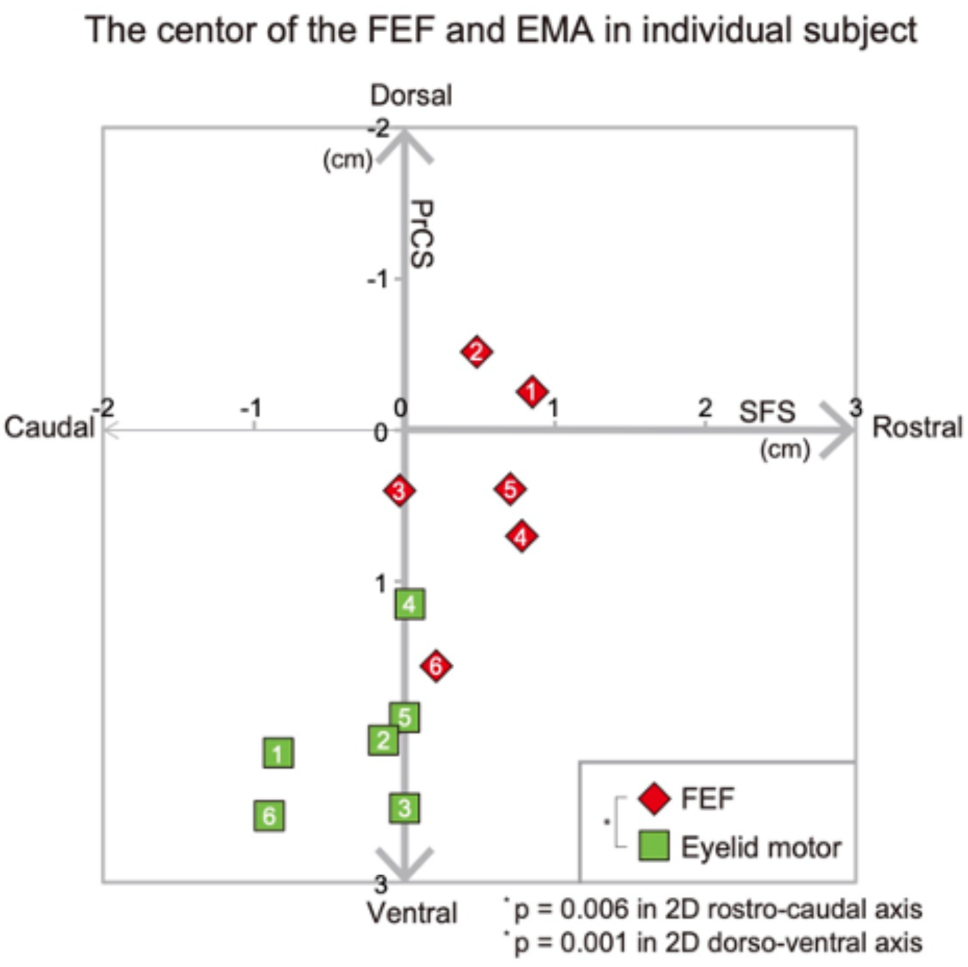
Centers of the FEF (argyle) or the EMA (square) electrodes in each subject in the study are shown. The number within each mark denotes the same subject.

### 3.6 Surrounding functions of the FEF and the EMA in 2D analysis

To investigate the location of the FEF in relation to the precentral motor cortex or the motor homunculus, the function of the region surrounding the FEF electrode was investigated (Fig. 5a). After eliminating both indeterminate electrodes and the FEF itself from all 52 electrodes, the number of remaining electrodes was 34 in the rostral region, 26 in the dorsal region, 38 in the ventral region, and 40 in the caudal region. Rostrally, the silent area was found almost exclusively (90%: 30.5/34 electrodes), whereas the face motor area was seen in a very small region (4%: 1.5/34 electrodes). Dorsally, the motor area was seen in 58% (15.5/26 electrodes) of FEF electrodes, the majority of which were found in the upper extremity motor area (54%: 14/26 electrodes). Ventrally, the motor area occupied a smaller region (38%: 14.5/38 electrodes) with the face motor area on the top (29%: 11/38 electrodes). Caudally, the FEF in all subjects was attached to the precentral motor cortex, mostly to the hand motor area (70%: 28/40 electrodes). Based on these results, it is highly likely that the FEF is either attached to or partially embedded into the precentral motor cortex.

**Figure 5.**
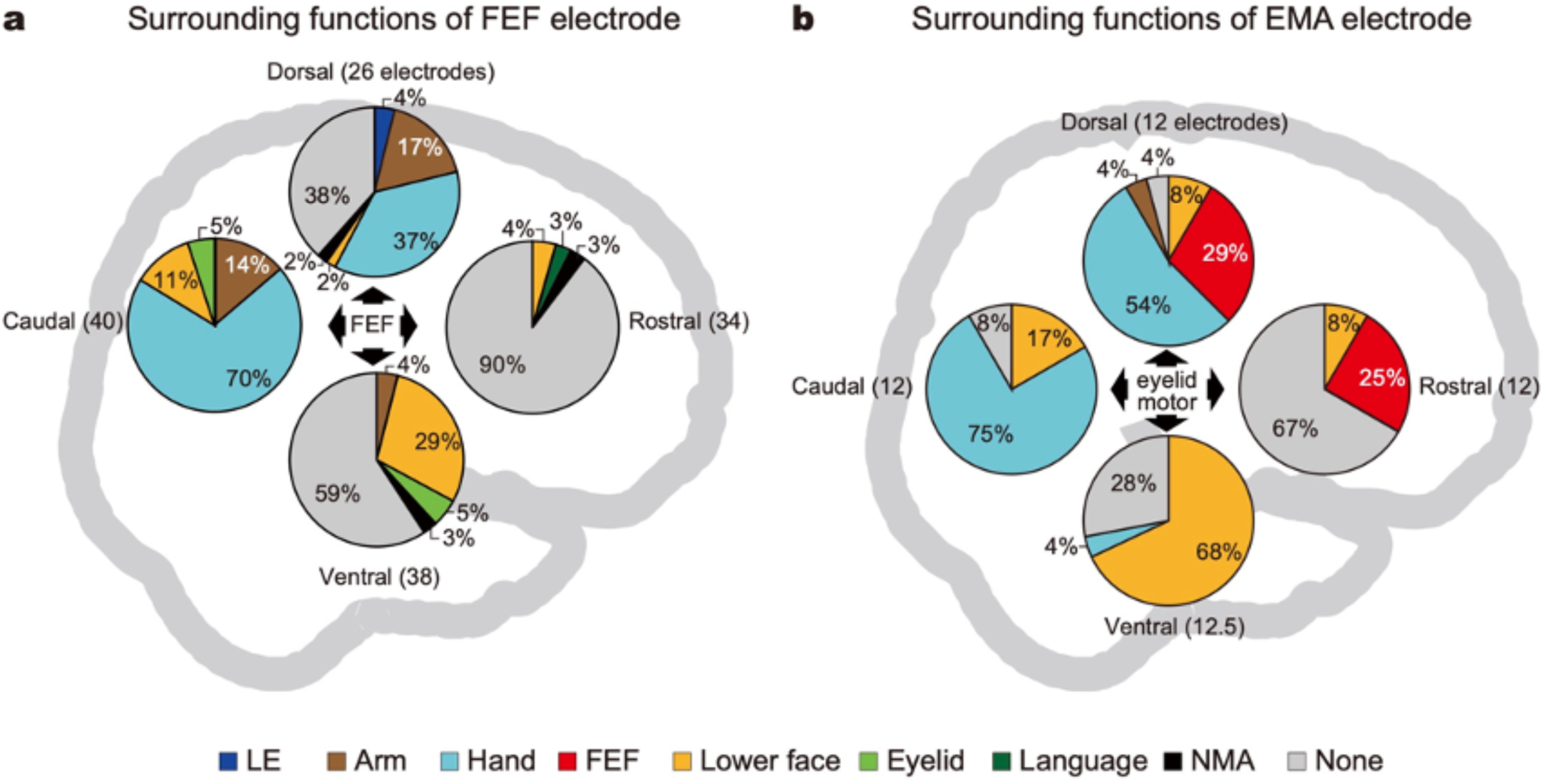
The ratio of each function surrounding the FEF (a) and (b) eyelid motor area.

The function of the region surrounding the EMA electrode was also investigated (Fig. 5b). After eliminating both indeterminate functions and those of the eyelid itself from all 14 electrodes, the number of remaining electrodes was 12 in the rostral region, 11.5 in the dorsal region, 12.5 in the ventral region, and 11 in the caudal region. Rostrally, while the silent area was mainly found (67%: 8/12 electrodes), the FEF was seen in a quarter of the region (25%: 3/12 electrodes). Dorsally, the most adjacent function was the hand (57%: 6.5/11.5 electrodes), followed by the FEF (30%: 3.5/11.5 electrodes). Ventrally, the lower face area occupied 68% of the region (8.5/12.5 electrodes). Caudally, the hand motor area was attached to three-quarters of the region (73%: 8/11 electrodes). Based on these results, the EMA was found embedded in the precentral motor cortex, situated caudal and ventral to the FEF. In the 2D dorso-ventral axis, the EMA was situated between the hand motor area and the lower face motor area.

The EMA localization within the precentral motor cortex is further confirmed by 3D analysis. Regarding the Z-axis, the EMA coordinates in the MNI standard space also showed a significant difference in distribution compared with all other positive motor areas. The EMA was situated significantly ventral to the distributions of LE**, arm**, forearm**, hand*, FEF*, and dorsal to lower face** and tongue** (Dunnett test: *p < 0.05, **p < 0.001) (Fig. 6).

**Figure 6.**
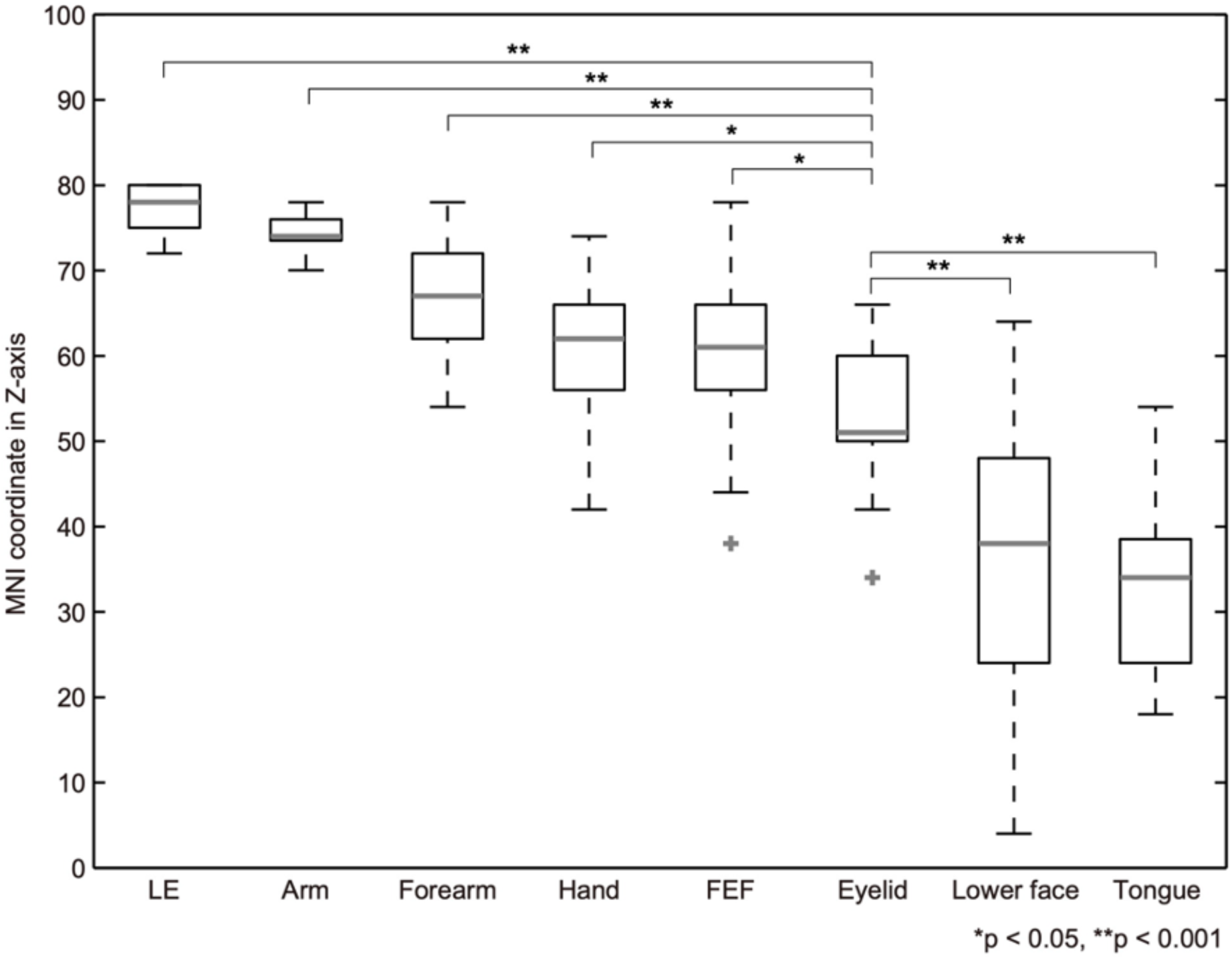
Boxplot representing the MNI coordinates in the Z-axis of each positive motor area. Asterisks (*, **) show significant distributional differences between the eyelid and other positive motor areas. The top and bottom portions of the box show the upper and lower quartiles, respectively. The median corresponds to the thick gray line within the box. Lines extend from the box to the maximum and minimum values of the data. Each cross represents an outlier within each positive motor area.

### 3.7 Distribution of the FEF, EMA, and other motor areas in the MNI space

In the MNI standard space, all coordinates with respect to each motor function are shown as spheres in Fig. 7a. The location of each sphere in Fig. 7b shows the mean coordinate of each function. The coordinates obtained from neuroimaging studies and other stimulations (ECS or transcranial magnetic stimulation (TMS)) are shown in reference to the FEF and EMA mean coordinates in the present study (Figs. 8a and 8b. See discussion for details).

**Figure 7.**
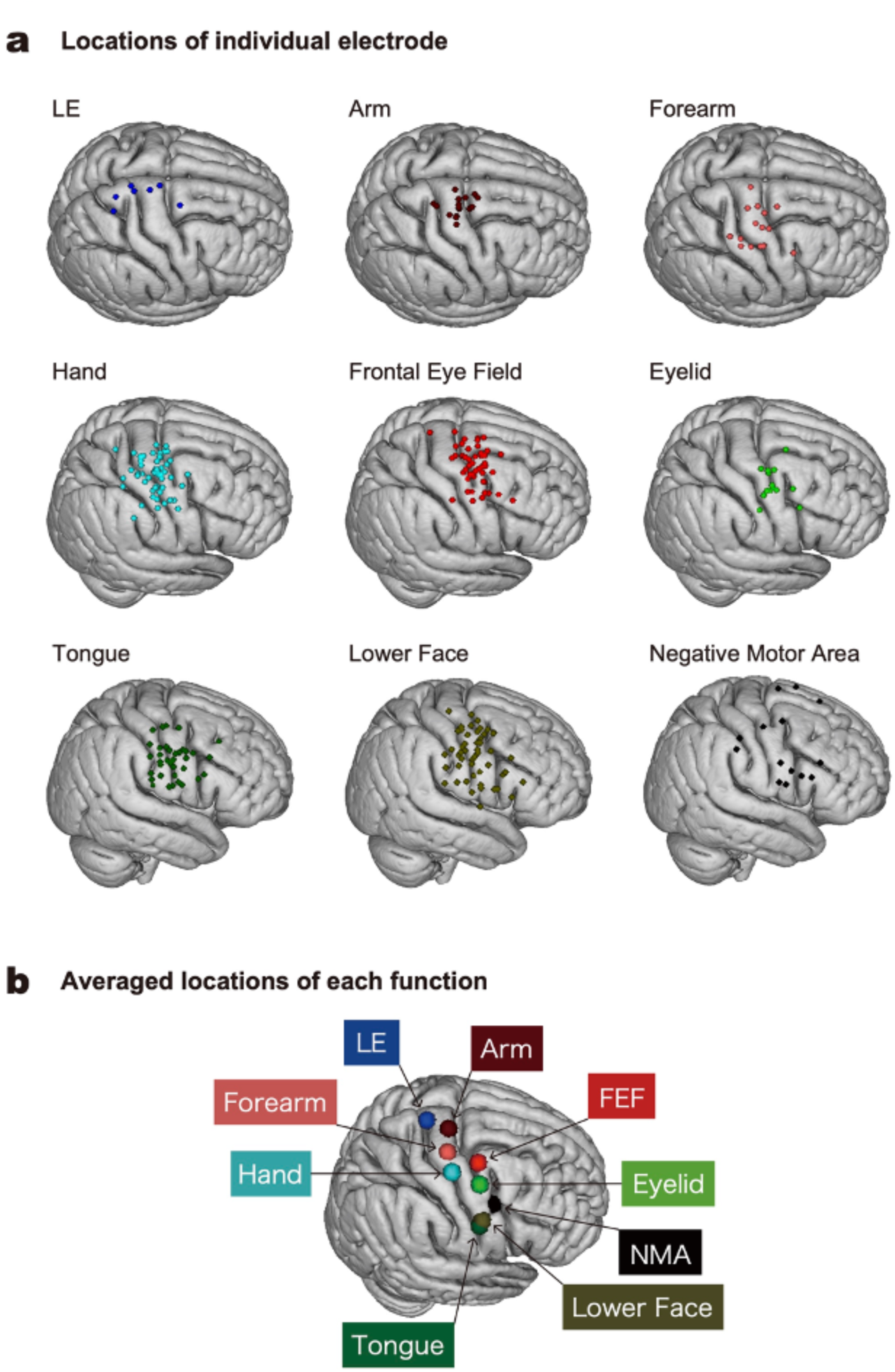
The localization map of each function in the MNI standard space. The location of each sphere shows co-registered (a) all coordinates from chronically implanted subdural electrodes and (b) averaged coordinates of each function.

**Figure 8.**
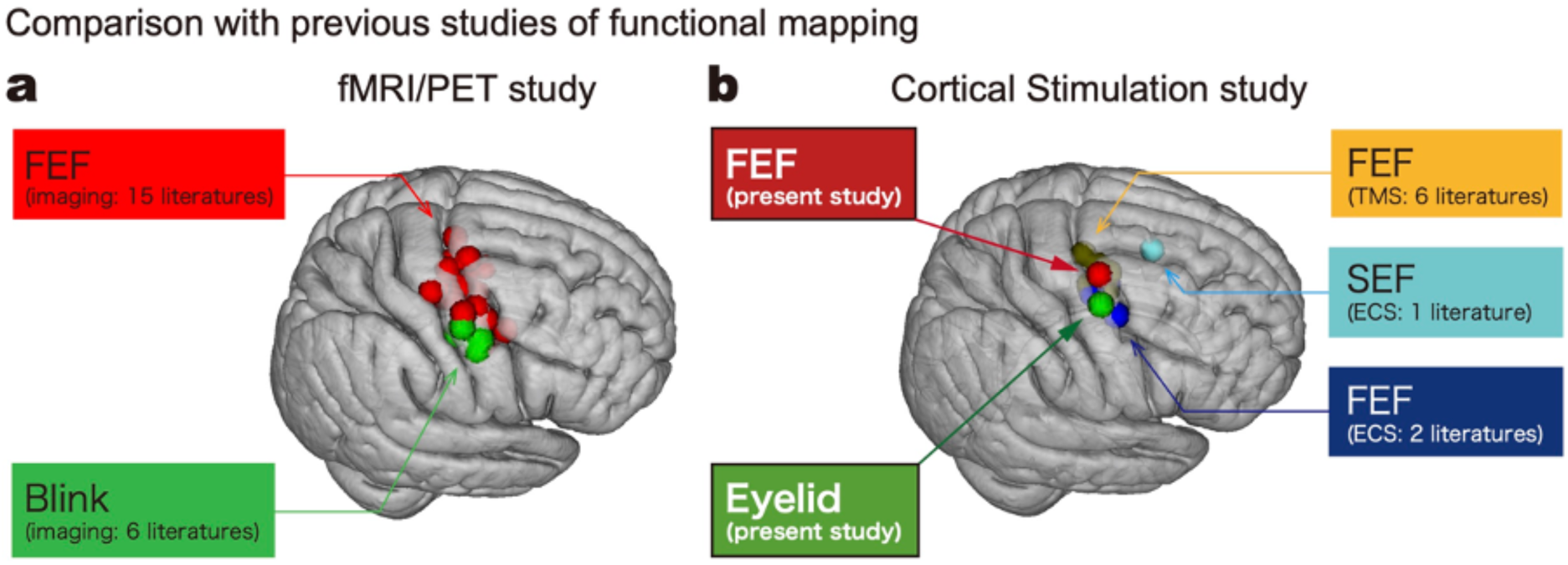
Averaged coordinates of each function in the MNI standard space (coordinates were flipped into the right hemisphere). (a) Comparison with previous functional neuroimaging studies. 15 literatures of FEF: (Paus, 1996; Petit et al., 1997; Corbetta et al., 1998; Culham et al., 1998; Luna et al., 1998; Berman et al., 1999; Petit & Haxby, 1999; Lobel et al., 2001; Gagnon, O’Driscoll, Petrides, & Pike, 2002; Kato & Miyauchi, 2003b; Koyama et al., 2004; Grosbras, Laird, & Paus, 2005; Amiez, Kostopoulos, Champod, & Petrides, 2006; Ettinger et al., 2008; van Koningsbruggen et al., 2012); 6 literatures of Blink: (Kato et al., 2003a, b; Bristow et al., 2005; Hanakawa et al., 2008; Amiez & Petrides, 2009; van Koningsbruggen et al., 2012). (b) Comparison with previous ECS and TMS mapping studies. 6 literatures of FEF (TMS studies): (Grosbras & Paus, 2002; Gagnon, Paus, Grosbras, Pike, & O’Driscoll, 2006; Morishima et al., 2009; Heinen, Feredoes, Weiskopf, Ruff, & Driver, 2014; Chen, Jin, Xiang, Liu, & Yin, 2018; Mastropasqua, Dowsett, Dieterich, & Taylor, 2019); 1 literature of SEF (ECS study): (Lobel et al., 2001); 2 literatures of FEF (ECS studies): (Blanke et al., 2000; Lobel et al., 2001).

## 4. Discussion

### 4.1 Highlight of the present study

By combining ECS and post-grid implantation MRI to achieve the best functional and anatomical precision in patients who had large subdural grid coverage over the premotor and perirolandic areas, the present study shed light on the relationship between the specific functional characteristics of FEF and their anatomical locations in reference to the SFS and PrCS. The majority of oculomotor response elicited by FEF stimulation was conjugated, saccadic eye deviation contralateral to the side of ECS. Head turning and non-saccadic eye deviation more frequently occurred in the vicinity of the PrCS. Functionally, the FEF was situated at the level of the hand motor area, more dorsal than was described in Penfield’s motor homunculus. The site of FEF protrudes anteriorly from the precentral motor stip. Anatomically, the majority of the FEF electrodes were located at BA 6 in the most-caudal region of the MFG, and in the adjacent part of the SFS or PrCS. The FEF distribution found in the present study supports the FEF localization provided by recent studies with both electrical stimulation (Blanke et al., 2000; Lobel et al., 2001) and TMS (Grosbras et al., 2002; Gagnon et al., 2006; Morishima et al., 2009; Heinen et al., 2014; Chen et al., 2018; Mastropasqua et al., 2019), as shown in Fig. 8b.

ECS of the EMA elicited eyelid closure and upper eyelid elevation, which were stronger in the eyelid contralateral to the side of ECS. The EMA was embedded within the precentral motor cortex, ventro-caudal to the FEF. Since the study by Rasmussen and Penfield (1948), to our knowledge, this study is the first report of the locational discrimination between the FEF and EMA in the MNI standard space in a relatively large number of epilepsy patients by ECS mapping.

### 4.2 Methodological benefit

We only selected the patients whose epileptic foci were outside the precentral/premotor area. Using the ECS method established by Lüders (1987), we carefully monitored the ECoG and confirmed the absence of afterdischarges during ECS mapping and investigated physiological cortical functions. Over seventy years ago, Rasmussen and Penfield (1948) measured the distance from the sulci to the location of each electrode using photographs and sketches by the surgeon. As described above, in the present study, the location of each electrode was identified with MRI taken after electrode implantation. Each electrode was identified in 2D MR images using its void signal due to the property of platinum alloy. These methods minimized the effect of brain distortion by the chronic implantation of the subdural electrodes. In addition, electrodes were non-linearly co-registered to the patient MRI obtained before implantation, and then to the MNI standard space. These approaches provided accurate functional mapping in a standard space and defined the locations of the FEF, EMA, and other positive/negative motor areas in a systematic manner.

### 4.3 Anatomical distribution of the FEF

All electrodes where ECS with a low-current threshold (∼5 mA) elicited oculomotor response were located relatively confined and near the intersection of the SFS and the PrCS (Fig. 2a). In contrast, the middle- (5.5–10 mA) and high- (10.5–15 mA) current-threshold groups were relatively more dispersed. Lobel et al. (2001) found that the oculomotor area at the intersection of the SFS and the PrCS could be most easily elicited by electrical stimulation. Blanke et al. (2000) also reported that electrodes with higher current thresholds were localized in the anterior and lateral parts of the oculomotor area. The findings of our study are consistent in the identification of low-current thresholds concentrated at the intersection of the SFS and PrCS. The distributions of three different current-threshold groups suggest that the oculomotor function is located superficially at the intersection, and deeply at its peripheral regions. A higher current intensity is necessary to deliver the current to the FEF at deeper sulcal regions of the PrCS or SFS or, alternatively, the threshold is indeed higher at the crown part of the cortex in the peripheral regions.

The distribution of the FEF electrodes revealed a longer FEF distribution along the 2D dorso-ventral axis than the 2D rostro-caudal axis in both inter-subject and within-subject analyses. A group of coordinates identified as FEF by eye saccade motion from individual neuroimaging studies (Paus, 1996; Petit et al., 1997; Corbetta et al., 1998; Culham et al., 1998; Luna et al., 1998; Berman et al., 1999; Petit et al., 1999; Lobel et al., 2001; Gagnon et al., 2002; Kato et al., 2003b; Koyama et al., 2004; Grosbras et al., 2005; Amiez et al., 2006; Ettinger et al., 2008; van Koningsbruggen et al., 2012) is summarized in Figure 8. This distribution follows the PrCS and is similar to the distribution of center coordinates of the FEF obtained from individual patients in this study. Most of the FEF is located in BA6, the agranular cortex, possibly accounting for the large variance in FEF distribution along the PrCS.

On the other hand, in the present ECS study, individual FEF electrodes in the whole group analysis and the centers of FEF electrodes in the individual patient analysis both showed more caudal distributions along the SFS as the FEF shifted more ventrally along the PrCS (Fig. 2b). These analyses suggested that the distribution of the FEF is not only limited to the sulcal part of the gyrus, namely the PrCS as neuroimaging studies reported but is also present in the crown part of the most-caudal MFG region, caudal SFG, and ventral PrCG.

### 4.4 Eye movement characteristics of the FEF

Three-quarters of the oculomotor response showed saccadic movement (Fig. 3a). The proportions of saccadic and non-saccadic movements were almost consistent with those reported in previous ECS studies (Godoy et al., 1990; Blanke et al., 2000). A previous fMRI study also suggested that the pursuit areas were more limited than the saccade areas (Petit et al., 1999). In the present study, all non-saccadic electrodes were concentrated within ± 0.5 cm of the PrCS, and more caudal to the saccadic electrodes (Fig. 3a). In a previous stimulation study of the primate FEF, Gottlieb (1993) found that pursuit was elicited in the fundus and posterior bank, while the saccade was elicited in the anterior bank of the arcuate sulcus. Though the number of investigated subjects was limited, saccade areas were located anterior to pursuit areas in both ECS (Blanke et al., 2000) and fMRI (Rosano et al., 2002) studies. In the present study, we were able to demonstrate distributional differences between saccadic and non-saccadic movements in a relatively large group of patients.

Recent studies demonstrated vertical eye movements by direct electrical stimulation with depth electrodes (Kaiboriboon, Lüders, Miller, & Leigh, 2012; Montemurro, Herbet, & Duffau, 2016). They suggested that the region involved in vertical eye movement is located at the deep and most-caudal part of the FEF. Here, we found that two-thirds showed purely horizontal eye movement and one-third showed oblique eye movement (Fig. 3b). The distribution of oblique eye movement extended more proximal to the PrCS compared with horizontal eye movement. These findings indicated that vertical eye movement is attributed to the electrical stimuli that affect relatively deep FEF regions situated at the sulcal part, namely, the PrCS.

One electrode showed disconjugate conversive horizontal eye movements located in the precentral gyrus at one of the most ventral positions among the FEF electrodes. Thurtell et al. (2009) also found that the stimulation of the electrode situated at the caudal end of the MFG resulted in conversive eye movement. Other nearby electrodes did not show such characteristics. Not only the electrode’s location but also other stimulus conditions such as the intensity of the electric current, individual variability, or electric current spreading via inter-cortical connections might be responsible for this rare symptom.

In the present study, one electrode over the PrCS showed HT without oculomotor response (Fig. 3c). Because the subject was able to keep his eyes fixated on an object during HT, this behavior is thought to be HT or unilateral neck motor response, which was dissociated from FEF or eye movement. This finding suggested that the neck motor area was located at the PrCS. However, HT without oculomotor response is found in a single electrode whereas no such behavior was elicited in nearby electrodes. As a plausible explanation of this finding, present ECS mapping might have had difficulties delivering a stimulation into the neck motor area and FEF separately. The neck motor area should be very limited whereas the FEF is widely distributed and located close to the neck motor area. The diameter of 3.97 mm of the subdural electrode we used would be too large to discriminate between these two functional areas. The stimulation over the PrCS might have the advantage of delivering the electrical stimuli to the neck motor area, which is likely situated at the sulcal part of the gyrus, the PrCS. This speculation is supported by the distributional relationship of HT and eye deviation (Fig. 3c). Five out of eight electrodes of HT before-end, where head turning started before the full eye deviation, were located over the PrCS. In contrast, only one out of fourteen electrodes was located over the PrCS, which showed eye deviation without HT. The different ratios implied the existence of the neck motor area at the PrCS.

In the dorso-ventral axis, the location of the HT before-end electrode was significantly dorsal to both HT after-end and eye deviation without HT electrodes (Fig. 3c). In contrast, every electrode of eye deviation without HT was located ventral to the SFS. This difference might stem from the three FEF electrodes with HT before-end situated dorsally in the SFG, which is associated with the supplementary eye field (SEF). In a study by Lobel et al. (2001), stimulation at the SEF resulted in contralateral HT at every electrode, whereas stimulation in the oculomotor area outside the SEF showed HT in only half of the electrodes. Based on the semiology of epileptic seizures, SEF in humans is thought to be closely associated with the head region in the SMA (Lim et al., 1994; Chee, 2000). Cortico-cortical connectivity is present between the SMA and the lateral premotor or precentral motor area (Matsumoto et al., 2007). HT before-end response in these three electrodes might have occurred since these electrodes were indeed the SEF itself or ECS of these electrodes led to the excitation of the SEF via cortico-cortical connections. Further ECS studies clarifying SEF characteristics of eye movements are needed to differentiate the SEF from the FEF.

### 4.5 Functional localization of the FEF

The FEF was attached to the motor cortex caudally in all patients, while in contrast, motor responses were almost exclusively absent in the rostral region (Fig. 5a). This means that the site of the FEF protrudes anteriorly from the precentral motor stip. The FEF was located between the upper extremity (in dorsal) and the lower face (in ventral). Caudal to the FEF, the majority of electrodes belonged to the hand motor area. These findings were consistent with Godoy’s ECS study (1990), which could not provide anatomical localization due to the lack of neuroimaging investigation. Based on this finding, our study suggests the FEF was attached to or partially embedded into the precentral motor cortex (Figs. 5a, 7b), more dorsal (Foerster, 1931; Penfield et al., 1954) than the location reported in previous stimulation studies. The functional location of the FEF is further supported by epicortical recordings of the motor-related cortical potential with saccade eye movements. The potential was recorded 1–2 cm rostral to the hand, arm, or face primary motor area (Yamamoto et al., 2004).

### 4.6 The eyelid movement characteristics of the EMA

Previous studies have suggested that blinking involves different regions of the brain depending on the nature of blinking – whether unconscious or voluntary. In a monkey study, Gong et al. (2005) found that cortical afferents into the orbicularis oculi motor neurons originated from multiple motor areas, including the lateral premotor areas and M1, not only from the FEF. In humans, activity in the medial frontal gyrus has been related to spontaneous or unconscious blinking, and activity in the precentral motor to voluntary blinking (Yoon, Chung, Song, & Park, 2005). Most fMRI studies investigated the functional brain regions responsible for voluntary blinking to define the EMA (Kato et al., 2003a; Bristow et al., 2005; Hanakawa et al., 2008; van Koningsbruggen et al., 2012).

In the present study, we investigated a “compulsory” eyelid motor response by electrical stimulation. Muscle contraction of either the orbitalis oculi (eyelid closure) or the levator palpebrae superioris muscle (upper eyelid elevation) was observed and stronger in the eyelid contralateral to the side of ECS. These are clearly different from the brief, bilateral blinks that we unconsciously perform in daily life, and are considered to be more like intentional eyelid movements. From the observed responses by ECS, it is likely that the EMA account for the voluntary control of the orbicularis oculi and levator palpebrae superioris muscles, which is bilateral but stronger on the contralateral side.

### 4.7 Anatomical and functional localization of the EMA v

To our knowledge, we are the first to functionally and anatomically identify the EMA by combining ECS and post-grid implantation MRI for precise localization of the electrodes. No such similar work has occurred since Penfield’s research in the mid 20^th^ century. Anatomically, the EMA was mainly located in the anterior part of the PrCG (Figs. 4, 7).

We also found that the EMA had a different location from the FEF, situated caudal and ventral to the FEF (Fig. 4). For subjects showing both eyelid motor and oculomotor response separately, plotting the centers of these motor areas showed clearly different coordinates in the 2D dorso-ventral axis and the 2D rostro-caudal axis, respectively (Fig. 4). These positional relationships between the eyelid motor and oculomotor responses were in line with the findings of a previous imaging study (van Koningsbruggen et al., 2012). In the 2D analysis with the Talairach Client software program, most FEF electrodes were localized rather rostral to the PrCS, while most EMA electrodes were localized rather caudal to the PrCS. On the other hand, in the 3D MNI standard space, there were no significant distributional differences in the Y-axis [rostro-caudal]. This discrepancy suggested that the distributional difference between the EMA and the FEF in the 2D rostro-caudal axis is based on the sulcal pattern of the PrCS. In fact, the PrCS itself runs more rostral as it proceeds to the ventral side of the cerebral convexity.

With regard to the surrounding functions, most of the EMA electrodes were adjacent to other positive motor areas, except for the rostral portion (Fig. 5b). The majority was constituted by the hand motor area dorsally and the lower face motor area ventrally. In contrast, most of the FEF is surrounded by the silent area, except for the caudal portion (Fig. 5a). This contrast is confirmatory evidence that the FEF is located anterior to the precentral motor cortex, while the EMA is embedded in the precentral motor cortex. Statistical analysis in the MNI standard space (the Z-axis) confirmed that the location of the EMA is significantly different from that of other positive motor areas in the motor homunculus, namely, the areas situated dorsal to the lower face and ventral to the FEF and the hand motor area (Fig. 6). This configuration further consolidates the anatomical localization for eye and eyelid movements reported in neuroimaging studies (Fig. 8a).

In a previous study carried out by Rasmussen and Penfield (1948), distributions of the FEF and EMA were not separate; both distributions ranged from the precentral gyrus immediately anterior to the central sulcus to the caudal portion of the frontal convolutions. They also reported that accurate localization in the superior, middle, or IFG could be uncertain because of the variability in human convolutional patterns and the limited exposure of the cortex during surgery. Compared with their method, namely, visual inspection during surgery, the present study had the advantage of positional identification of electrodes, which may contribute to a relatively low variance in each function and show different distributions in the FEF and the EMA.

### 4.8 Motor homunculus in the precentral motor cortex

The localization maps for each motor function were identified in the MNI standard space. The mean coordinates of LE, arm, forearm, hand, FEF, EMA, lower face, and tongue were identified in dorsal to ventral order, mainly in the PrCG (Fig. 7b). All of these mean coordinates were located in BA6, in the anterior part of the PrCG, including that of the FEF. NMA electrodes were discretely placed throughout the premotor cortex as in previous ECS studies (Mikuni et al., 2006; Borggraefe et al., 2016). The mean coordinate of the NMA was located in BA6, over the PrCS.

Anatomically, the EMA was localized to the anterior part of the precentral gyrus and functionally embedded in the precentral motor cortex between the hand and lower face motor areas (Fig. 6). The location of the EMA and the surrounding functional areas were consistent with the results of previous fMRI studies using voluntary blink movements (shown in Figs. 7 and 8b). To our knowledge, since Penfield’s research, this is the first study to functionally and anatomically identify the EMA using ECS.

### 4.9 Limitations

Direct electrical stimulation of the human cortex is the gold standard for mapping brain functions. However, this study had several limitations. First, the location and size of the subdural grid electrodes were guided by a clinical purpose and not based on scientific criteria. Because some FEF electrodes were placed at the edge of the grid or located in a cluster with other FEF electrodes, the analysis of functions surrounding the FEF was partly compromised. Second, the stimulation intensity was started under-threshold. The stimulation intensity and duration were carefully increased to the first point that showed a positive/negative motor response in each electrode. However, we cannot deny the possibility that cortical activation indirectly via the intracortical white matter pathway is ultimately causally associated with the function of the FEF. Third, we only analyzed the location of the crown region and not the sulcal region to complement fMRI studies that highlight activation in the sulcal region. In addition, recent ECS studies with depth electrodes found that the stimulation of the deep PrCS or white matter tracks underneath the FEF is also involved in the control of eye movement (Kaiboriboon et al., 2012; Montemurro et al., 2016). Investigating the sulcal region and subcortical structures with SEEG in a large group of patients would provide further findings regarding the functions and roles of the FEF.

## 5. Conclusions

The present combined ECS and MRI study confirmed that the FEF is anatomically distributed from the intersection of the SFS and the PrCS to the most-caudal region of the MFG, and is located rostral to the hand motor area. The motor characteristics elicited by stimulation of the FEF, such as eye movements (saccades or non-saccades, etc.), the direction of head rotation (horizontal or oblique), and the order of these movements occur, differed depending on the distance from the stimulation site to the PrCS. The stimulation of the ventral and caudal sides of the FEF elicited eyelid movements such as eye closure and upper eyelid elevation. Eyelid muscle contraction was stronger on the opposite side of the stimulus. This is the first report of the EMA identified by ECS since the work of Penfield. A standardized map of the FEF, EMA, and precentral motor homunculus is provided for reference to human system neuroscience research.

## Acknowledgments

We wish to thank Timothy O’Connor, Karl Horning, and Mary Jo Sullivan for their technical assistance, and Eric LaPresto for constructing an in-house image rendering software.

## Author Contributions

**Tomoyuki Fumuro:** Investigation, Formal analysis, Writing-Original draft, Visualization, **Riki Matsumoto:** Conceptualization, Investigation, Formal analysis, Data Curation, Resource, Methodology, Supervision, Writing-Review & Editing, Funding Acquisition, **Juan Bulacio**: Data Curation, Resource, Methodology, **William Bingaman:** Data Curation, Resources, Methodology, **Akio Ikeda:** Validation, **Hiroshi Shibasaki:** Supervision, **Hans O. Lüders:** Conceptualization, Resource, Methodology, Project administration, **Dileep R. Nair:** Conceptualization, Investigation, Formal analysis, Data Curation, Resource, Methodology, Writing-Original draft, Project administration

## Ethics

This study was approved by the Institutional Review Board Committee of the Cleveland Clinic Foundation (IRB #4513). All procedures were conducted in accordance with the Declaration of Helsinki. Informed consent was obtained from all participants or their legal guardians.

## Data and Code Availability

The data that support the findings of this study are available from the corresponding author upon reasonable request.

## Abbreviations

ANOVA: analysis of variance
BA: Brodmann’s area
ECoG: electrocorticogram
ECS: electrical cortical stimulation
EMA: eyelid motor area
FEF: frontal eye field
FSL: FMRIB Software Library
HT: head turning
LE: lower extremity
MANOVA: multivariate analysis of variance
MFG: middle frontal gyrus
MNI: Montreal Neurological Institute
PrCG: precentral gyrus
PrCS: precentral sulcus
SEEG: stereotactic-electroencephalography
SFS: superior frontal sulcus

